# Age-dependent peripheral nerve and Schwann cell abnormalities in a mouse model of late-onset spinal muscular atrophy

**DOI:** 10.64898/2026.07.27.740920

**Authors:** Kai Christine Liebig, Nicola Bense, Linda-Isabell Schmitt, Stefanie Hezel, Christoph Kleinschnitz, Markus Leo, Tim Hagenacker

**Affiliations:** Department of Neurology, Center for Translational and Behavioral Sciences (C-TNBS), University Medicine Essen, Essen, Germany

**Keywords:** SMA, SC, neuroinflammation, mouse model, macrophages, Sox2, Sox10, g-ratio, myelination, peripheral nerve

## Abstract

Spinal muscular atrophy (SMA) is increasingly recognized as a multisystem disorder involving non-neuronal cells, yet the role of Schwann cells (SCs) in late-onset SMA (loSMA) remains unclear. We investigated age-dependent peripheral nerve pathology in a four-copy *SMN2* mouse model of loSMA. Sciatic nerves from wild-type and loSMA mice were analyzed at postnatal (P) days 20, 35, 70, and >100 using semi-thin morphometry, immunofluorescence for MBP, Sox10, Sox2, and F4/80, and nerve conduction studies. loSMA nerves showed reduced myelin thickness at all time points and smaller axon diameters at P20 and P35. G-ratios were reduced at P20 but increased from P35 onward, indicating progressively altered axon-myelin relationships. MBP immunofluorescence intensity, compound muscle action potential amplitude, and nerve conduction velocity were reduced in loSMA mice at P>100. The proportion of Sox2^+^ SCs increased from P35 onward, while Sox10^+^ cell abundance increased at later stages. F4/80^+^ macrophages were transiently elevated at P35 and correlated with Sox2^+^ cell numbers at this stage. These findings demonstrate age-dependent myelin abnormalities, altered SC states, and transient accumulation of macrophages in loSMA peripheral nerves. Whether these changes are SC-autonomous or secondary to chronic axonal dysfunction remains to be determined.

## Introduction

Spinal Muscular Atrophy (SMA) is a hereditary disease characterized by degeneration of spinal motor neurons (MNs), leading to severe disability. The loss of MNs is caused by a mutation in the *survival of motor neuron 1 (SMN1)* gene, rendering the resulting SMN protein nonfunctional. Only low levels of functional SMN protein are produced from the *SMN2* gene, resulting in heterogeneous clinical phenotypes of SMA^1–3^. The existing SMN-enhancing therapeutics are efficient at preventing motor neuron loss in presyptomatic SMA patients, but therapeutic results vary strongly in symptomatic patients especially in long-standing late-onset SMA (loSMA) ^4–6^. The limited amelioration of symptoms observed in many loSMA patients indicates additional pathomechanisms affecting other cells or tissues beyond spinal MNs. SMA-related pathological changes have been shown in various other organs ^7^ ^8^. These observations highlight the unmet need for additional therapeutic options targeting other underlying pathomechanisms.

Glial pathology contributes substantially to SMA ^9–11^. In the four-copy SMN2 mouse model, astrocyte activation precedes motor neuron loss, and modulation of astrocyte-associated pathways can reduce motor neuron stress ^12–14^. The comparatively slow and staged disease course of this model provides an opportunity to investigate peripheral cellular changes that may emerge before or during progressive motor decline ^15^. Schwann cells (SCs) are essential for peripheral axon support, myelination, and nerve conduction. Studies in severe early-onset SMA models have demonstrated intrinsic SMN-dependent SC abnormalities and partial functional improvement following SC-specific SMN restoration ^16, 17^. Dysfunction of SCs leads to various pathological changes, including demyelination, axonal degeneration, and neuroinflammation ^18^. In loSMA, SC injury could arise either as a primary consequence of SMN deficiency or secondarily in response to chronic axonal dysfunction. In either case, impaired myelination, axonal support, or macrophage-associated nerve remodelling could further compromise peripheral conduction and motor function.

The contribution of Schwann cells and peripheral nerve pathology to loSMA has not yet been defined. We therefore characterized age-dependent changes in sciatic nerve morphology, myelin-associated MBP immunofluorescence, SOX10- and SOX2-positive Schwann cell populations, F4/80-positive macrophages, and electrophysiological function in the four-copy SMN2 mouse model. We aimed to determine whether structural and cellular abnormalities emerge before late-stage functional impairment and whether Schwann cell-associated changes are accompanied by macrophage accumulation.

## Methods

### Animal model

SMN-deficient mice (FVB.Cg-*SMN1*^tm1Hung^Tg(*SMN2*)2Hung/J; loSMA mice) were purchased from Jackson Laboratory (#005058, Bar Habor, ME, United States). These mice were homozygote for the murine *SMN1* knockout and the insert of human *SMN2* (four copies), reflecting human loSMA ^15, 19^.

Age-matched wild-type (WT) FVB/N mice served as control. LoSMA mice are smaller in size and have decreased motor functions with MN loss occurring from postnatal day (P) 35 onwards with preceding astrocyte activation ^13^. Therefore, the following time points were used here: P20 (spinal astrocyte activation), P35 (onset of MN loss), P70, and P>100 (later disease stages). Both sexes were included. Animals from different litters were used for each experiment and were randomly picked. The study was not powered to formally assess sex-dependent effects, and sex-stratified results are therefore reported descriptively.

The animals were kept on a 12/12-h light/dark cycle with water and standard food pellets available ad libitum. Animals were monitored weekly to examine body condition, weight, and general health.

Mice were maintained and bred in the Animal Research Lab of the University Medicine Essen. All experiments were conducted in accordance with the animal welfare guidelines of the University of Duisburg-Essen. The use of the late-onset SMA mouse model was approved by the State Office for Consumer Protection and Food Safety (LAVE), North Rhine-Westphalia,

Germany (approval number 81–02.04–2020.A335). The number of animals used for the experiments was in accordance with the 3Rs concept.

### Tissue preparation

Sciatic nerves of loSMA and WT mice were harvested at P20, 35, 70, and >100. Briefly, animals were anesthetized and sacrificed. Sciatic nerves were removed and processed by placing them into TissueTek tissue molds and freezing them in isopropanol on dry ice. Tissue was stored at −80° C until use.

### Nerve conduction studies

Electrophysiological experiments were conducted immediately after the animal was sacrificed. Recordings of the compound muscle action potential (CMAP) were used to identify changes within the motor nerve conduction. First, the sciatic nerve was stimulated at the sciatic notch using microneedle electrodes, then at the ankle. Finally, the reference electrode was placed at the nape of the neck. A single pulse was delivered each time using 5 Hz for 0.1 ms. The subtracted latency of both measurements, as well as the amplitude, was recorded.

### Semi-thin sections

Sciatic nerves were harvested and immediately fixed for 24 h using 4 % formaldehyde and 2.5 % glutaraldehyde in 0.1 M PHEM buffer. After fixation, nerves were washed three times with 0.1 M PHEM buffer. Nerves were further processed using a microwave for contrast and embedding (TedPella Biowave) and stained with 2% osmium tetroxide and 2% uranyl acetate. Afterward, the tissue was dehydrated and embedded in epoxy resin. Resin hardening was performed at 60 °C for 3 days.

Embedded nerves were cut in semi-thin cross sections (500 nm) on a Leica UC7 ultramicrotome, contrasted using toluidine blue and preserved with mounting medium.

For every time point five loSMA and WT mice were used. Images of semi-thin sections were acquired using a Leica microscope. Myelin sheaths were measured using ImageJ to obtain parameters like thickness, axon diameter and g-ratio.

### Immunostaining

Sciatic nerves were harvested, immediately embedded in mounting medium, and frozen. Using a Cryotome, 10 µm sections were cut longitudinally. Tissue preparation for fluorescence staining was performed as described elsewhere^13^. Nerves were stained with anti-Sox10 (#ab155279, Abcam), anti-Sox2 (#NBP3-31961, R&D Systems), anti-F4/80 (#MCA497G, BioRad), anti-SMI32 (#601712, Biolegend), anti-SMI35 (#835614, Biolegend) and anti-MBP (#PA1-10008, Thermo Fisher Scientific) antibodies and corresponding fluorescence labelled secondary antibodies as well as DAPI.

### Data analysis

Fluorescent images were acquired using a Leica Dmi8 microscope. Counts of Sox2^+^, Sox10^+,^ and F4/80^+^ cells were made with ImageJ. In short, images were converted to binary, thresholds were adjusted, and Sox2^+^ and Sox10^+^ cells were automatically counted. F4/80^+^ cells and DAPI signals were counted manually. Percentages of Sox10^+^ and Sox2^+^ cells in relation to DAPI counts and Sox2^+^ cells in relation to Sox10^+^ counts were calculated. For each nerve, the same area was used to count cells. For MBP, immunofluorescence signals were used to calculate corrected total cell fluorescence (CTCF), and CTCF was divided by the area size of the corresponding nerve. The CTCF/Area ratio was used for statistical analysis. In both cases, SMA and WT nerves were paired, and a ratio was calculated, which was logarithmically transformed and tested for significance using a one-sample ratio t-test against a hypothetical value of 1.

For the analysis of axon density shown in Fig. 2B, standardized mean differences between loSMA and WT mice were additionally calculated separately for each age using Cohen’s *d*. Cohen’s *d* was calculated as the difference between the group means divided by the pooled standard deviation, using one mean axon-density value per animal.

## Results

### LoSMA mice SC have thinner myelin sheaths and smaller axons

Semi-thin cross sections of sciatic nerves were analysed for changes of myelin sheaths in loSMA mice (Fig. 1a). Myelin sheath thickness of loSMA was reduced across all examined timepoints compared to WT mice (P20, P35, P70, P>100: *p*<0.001) (Fig. 1b). Meanwhile, axon diameter of loSMA mice was reduced at P20 and P35 (*p*<0.001) but not at P70 (*p*=0.59) and P>100 (*p*=0.87). Interestingly, violin plots show an inverse distribution of axon diameters, with a greater number of smaller (< 4 µm) and larger (> 6 µm) axons in loSMA than the mostly mid-sized axons in WT mice (Fig. 1c). G-ratio of loSMA mice was reduced at P20 (*p*=0.01) but elevated at all other timepoints (P35, P70, P>100: *p*<0.001) (Fig. 1d).

**Fig. 1.**
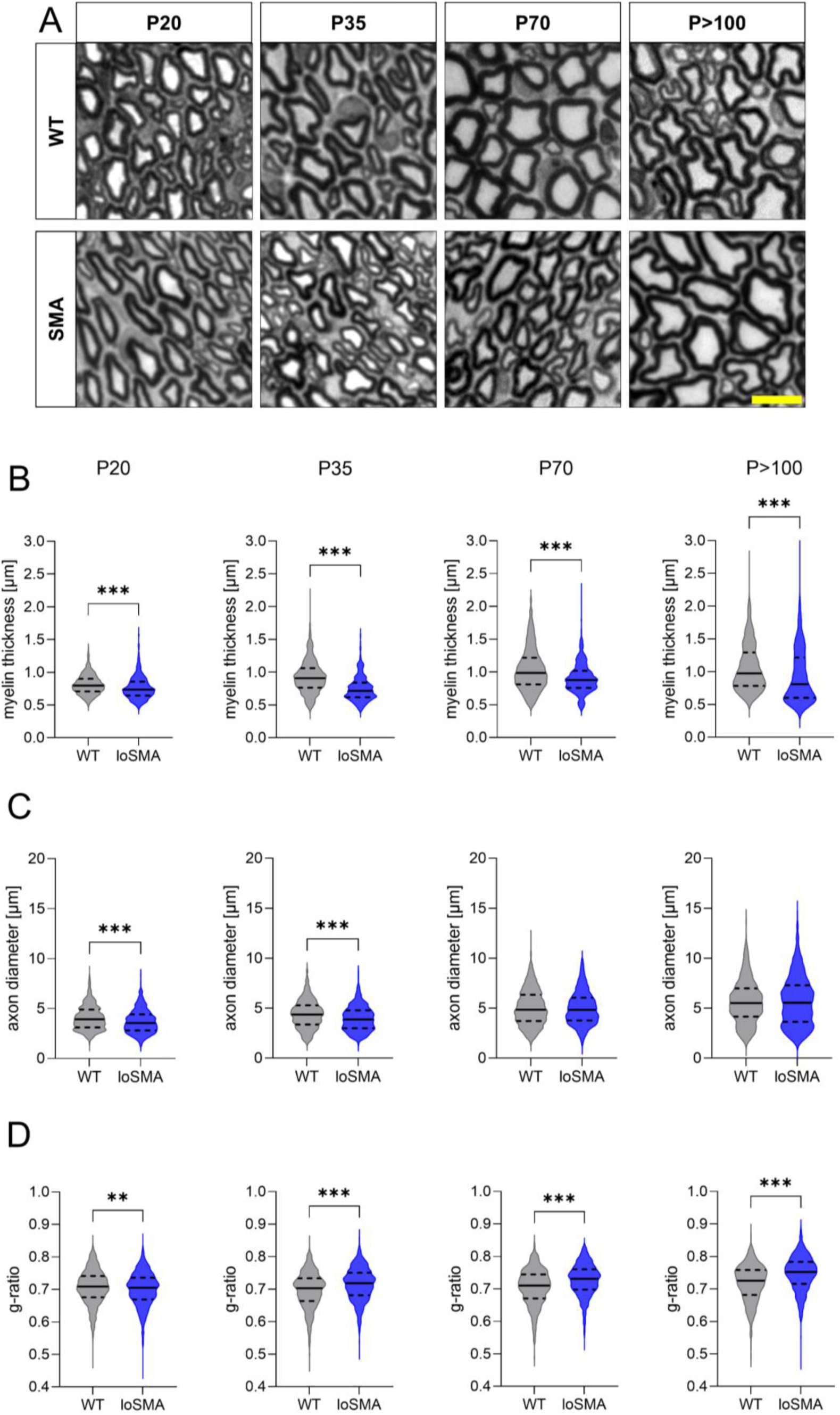
Semi-thin sections of sciatic nerves reveal smaller axons and thinner myelin sheaths in loSMA mice. (A) Representative images of semi-thin sections of WT and loSMA mice’s sciatic nerves, aged P20 to P>100, used for analysis. Scale bar equals 10 µm. (B) Myelin thickness was reduced in loSMA mice across all time points (P20, P35, P70, P>100: *p*<0.001). (C) In comparison, axon diameter was only significantly reduced at P20 and P35 (*p*<0.001). While there was no significant difference at P70 (*p*=0.59) and P>100 (*p*=0.87), violin plots showed an inverse distribution of axon diameters with a higher amount of smaller (< 4 µm) and larger (> 6 µm) axons in loSMA than the mostly mid-sized axons of WT mice. (D) G-ratio of loSMA mouse axons was elevated starting at P35 (P35, P70, P>100: *p*<0.001) but reduced at P20 (*p*=0.01). ns = not significant, * = *p*<0.05, ** = *p*<0.005, *** = *p*<0.001, statistical significance was analysed using Mann-Whitney Tests. Five animals of each condition and age were tested (n=5).

**Fig. 2.**
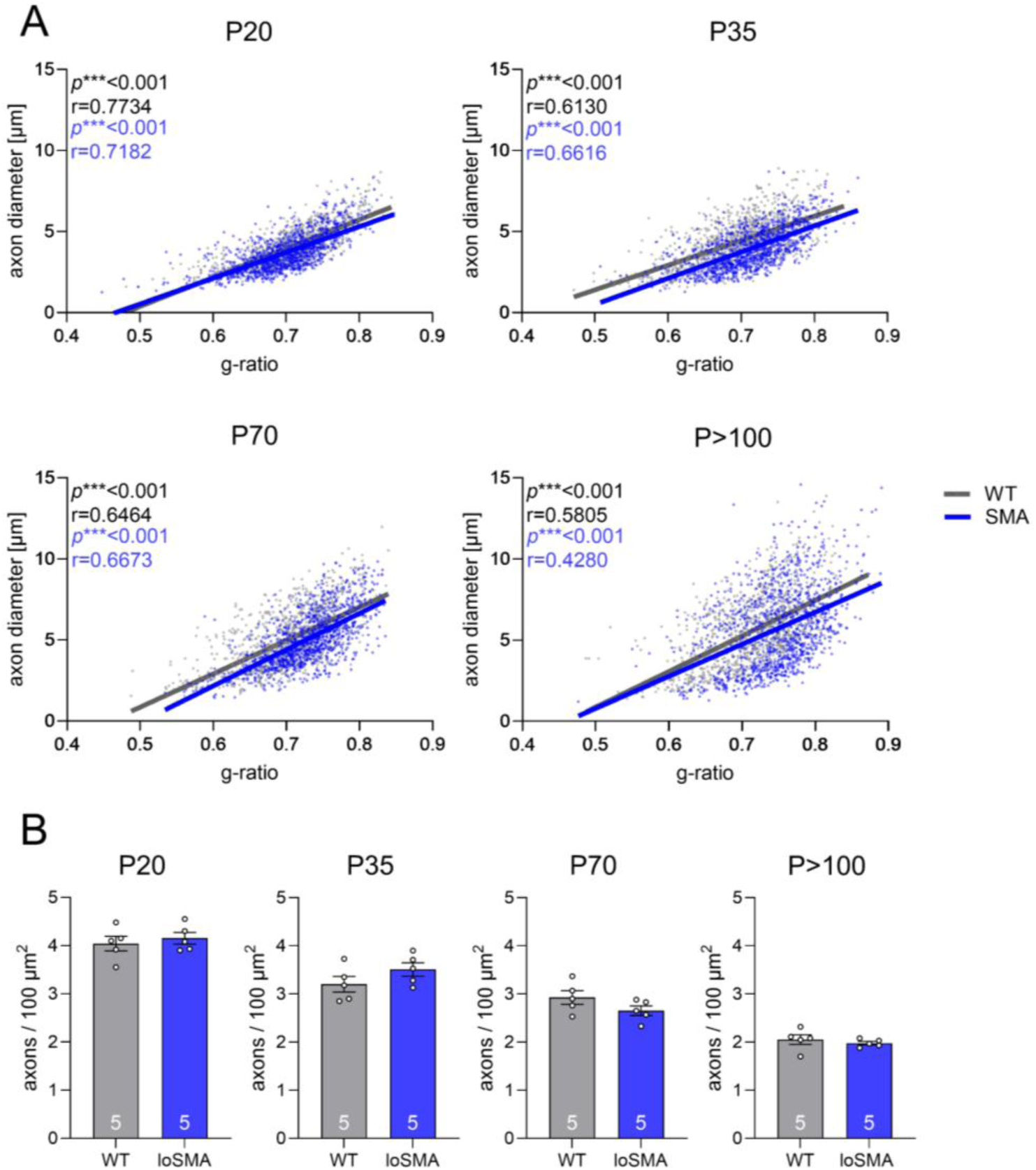
**Scatter plots of axon diameter versus g-ratio of loSMA (blue) and WT (grey) mice sciatic nerves across timepoints P20 to P>100 and number of axons per 100 µm**^2^. (A) Linear regression was similar between groups at P20 but showed a distinct shift towards smaller axons and an elevated g-ratio of loSMA mice from P35 onwards. Correlation of g-ratio and axon diameter was significant at all time points in both groups (P20, P35, P70, P>100: *p*<0.001). Statistical significance was analyzed using simple linear regression of the correlation matrix. (B) No significant difference was detected at any time point. Effect sizes for WT/loSMA comparison were small at P20 (Cohen’s d = 0.30), large at P35 (d = 0.90), moderate-to-large at P70 (d = 0.70), and moderate at P>100 (d = 0.60). Statistical significance was analyzed using an unpaired t-test. Five animals of each condition and age were tested (n=5).

These observations were confirmed by significant correlations of axon diameter and g-ratio (WT and loSMA axons across all time points; P20, P35, P70, P>100: *p*<0.001). While regression lines at P20 showed a similar trend, from P35 onwards a clear shift toward smaller axons with a larger g-ratio was observed (Fig. 2a).

Analysis of the number of axons per 100 µm^2^ revealed no significant differences between loSMA and WT mice nerves but indicated a trend to more axons at P20 and P35 and a slightly lower count at P70 and P>100 in loSMA mice. Therefore, effect sizes were calculated additionally. Effect sizes of the WT/loSMA comparison were small at P20 (Cohen’s *d* = 0.30), large at P35 (*d* = 0.90), moderate-to-large at P70 (*d* = 0.70), and moderate at P>100 (*d* = 0.60). (Fig. 2b).

### MBP levels are altered significantly in older loSMA mice

To identify possible changes in the amount of myelin between WT and loSMA mice, myelin protein MBP was measured via immunofluorescence in longitudinal sections of sciatic nerves. While no significant differences were detected at P20, P35, or P70, P>100 loSMA mice had lower MBP immunofluorescence intensity than WT mice (*p*<0.05) (Fig. 3).

**Fig. 3.**
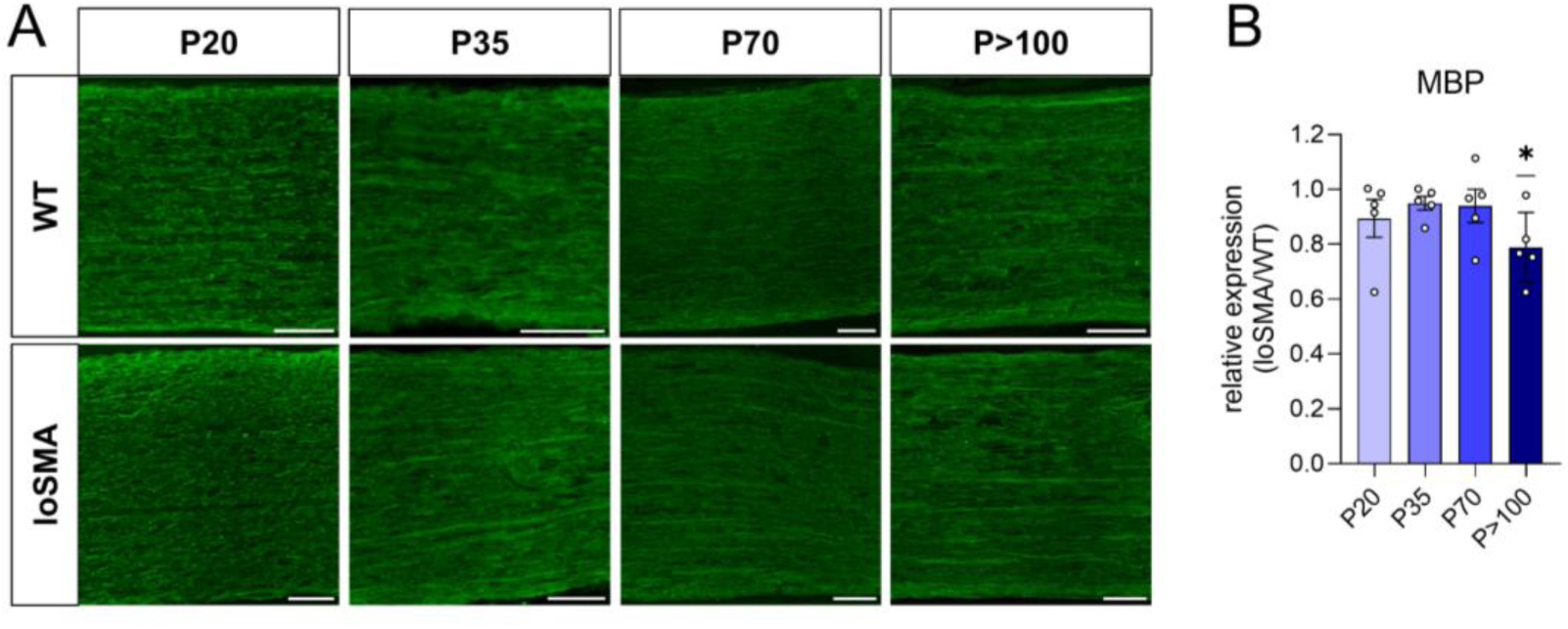
MBP immunofluorescence intensity is reduced in old loSMA mice. (A) Sciatic nerves were cut longitudinally and immunofluorescently stained for MBP (green). (B) The intensity of MBP was significantly reduced at P>100 in loSMA mice (*p*<0.05). * = *p*<0.05; statistical significance was analyzed using a one-sample ratio t-test by testing the logarithmic quotient of SMA/WT against a hypothetical value of 1. Five animals of each condition and age were tested (n=5). Scale bar equals 100 µm.

### Numbers of non-myelinating SC in SMA are increased

Myelinating and non-myelinating SCs express the transcription factor Sox10, but only non-myelinating SCs express Sox2. Immunostaining of longitudinal sciatic nerve sections of WT and loSMA mice showed a shift in abundance of Sox10^+^ and Sox2^+^ SCs. The overall number of Sox10^+^ SCs in comparison to all present cells (labeled using DAPI) was increased in loSMA mice at P70 and P>100 (\*\**p*<0.005), indicating an elevation of SC number. Furthermore, number of Sox2^+^ SCs was elevated in loSMA mice already at P20 (even though not statistically significant yet), reaching its peak at P35 and decreasing at P70 and P>100 (P35, P70, P>100 = *p*<0.05) both when compared to all present cells (labelled using DAPI) and only to Sox10^+^ SCs (P35, P70, P>100 = *p*<0.05) (Fig. 4).

**Fig. 4.**
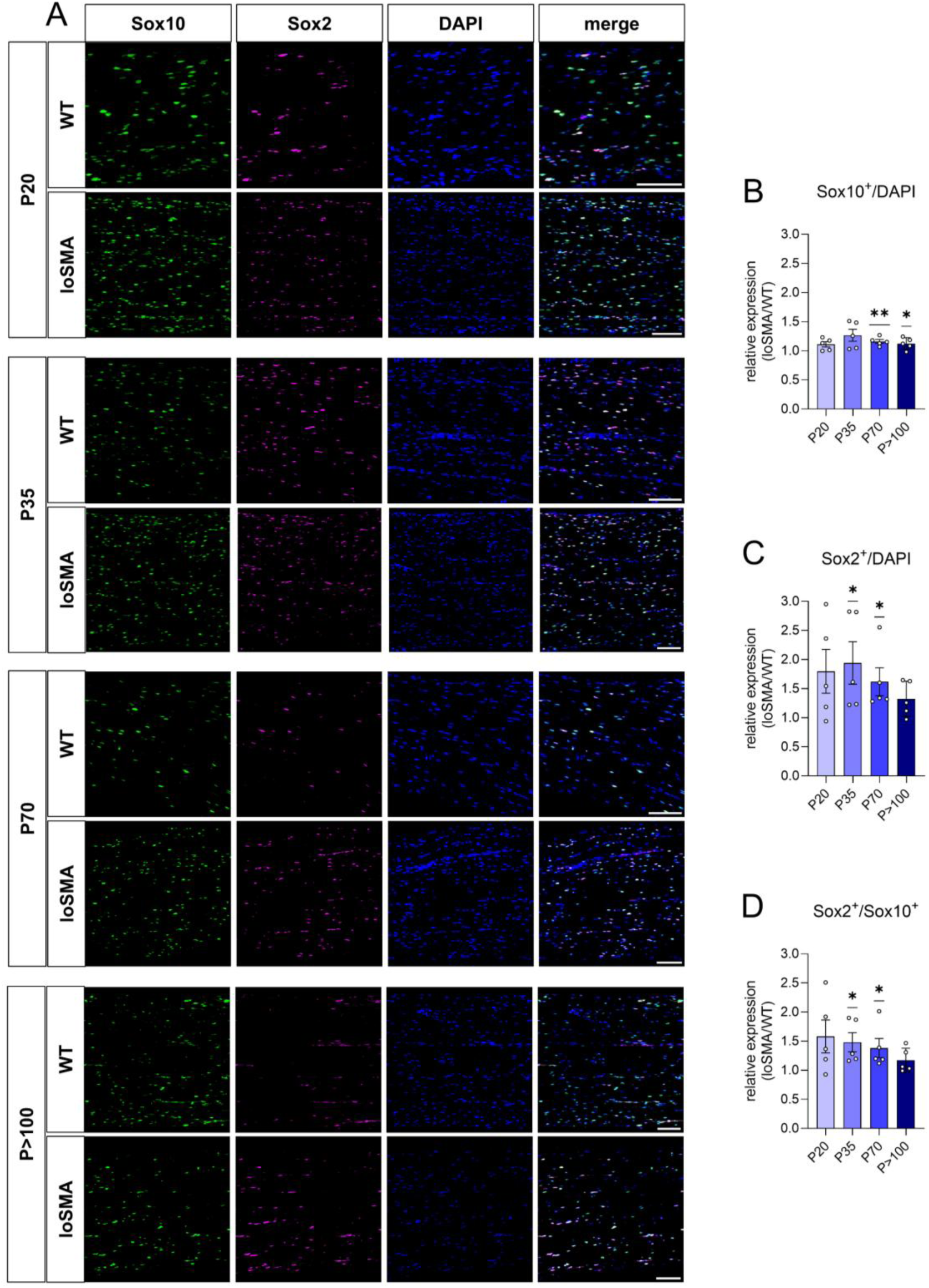
**Abundance of SCs analyzed using immunostaining is altered in longitudinal sciatic nerve sections of loSMA mice**. (A) Sciatic nerves of WT and loSMA mice were cut longitudinally and immunostained for SC markers Sox10 (green) and Sox2 (magenta) as well as DAPI (blue), images show a representative example of each time point. Merged images display an overlay of all channels. (B) The amount of Sox10^+^ SCs in comparison to all cells (labeled using DAPI) was significantly elevated from P70 onwards (P70, P>100: *p*<0.005), with a trend already showing at P35 (*p*=0.052). (C) Furthermore, the number of Sox2^+^ SCs was increased in loSMA mice at P35, P70 and P>100, reaching its peak at P35 (P35, P70, P>100 = *p*<0.05). (D) A similar observation was made comparing Sox2^+^ to Sox10^+^ cells (P35, P70, = *p*<0.05). * = *p*<0.05, ** = *p*<0.005, statistical significance was analysed using one sample ratio t-test by testing the logarithmic quotient of SMA/WT against a hypothetical value of 1. Five animals of each condition and age were tested (n=5). Scale bar equals 100 µm.

### Electrophysiological properties of loSMA sciatic nerves

Electrophysiological measurements of the sciatic nerves were performed to detect pathological changes in loSMA mice. Amplitude and nerve conduction velocity were on the same level in WT and loSMA mice at P35 but significantly decreased at P>100 in loSMA mice (P>100 = *p*<0.001, Fig. 5).

**Fig. 5.**
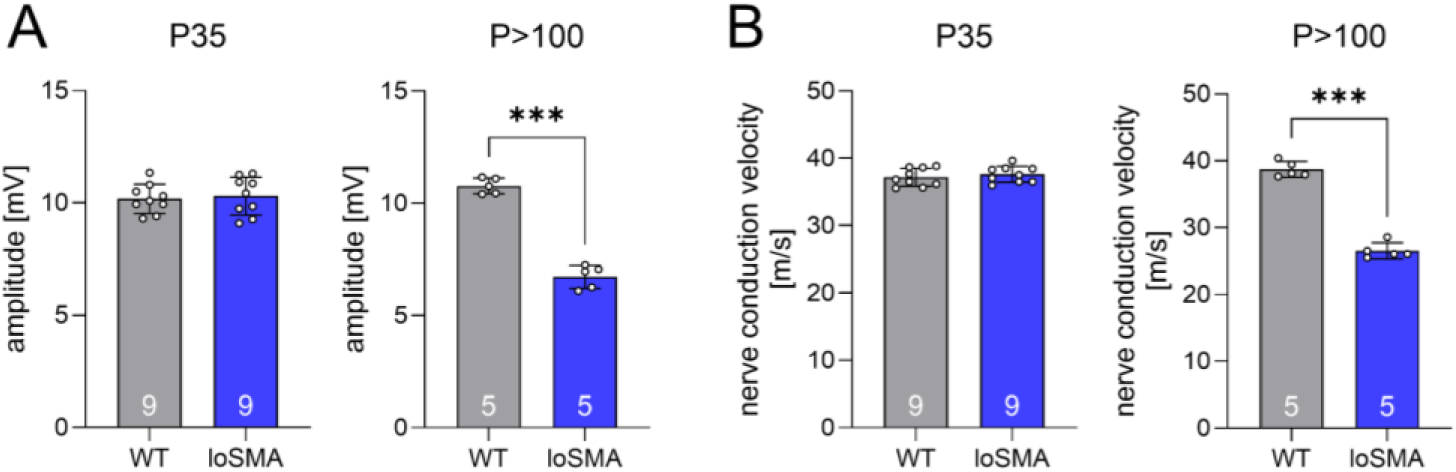
Nerve conduction studies on sciatic nerves revealed lower readings in old loSMA mice. (A) Amplitude was not altered at P35 between groups but decreased significantly in P>100 loSMA mice (blue, P>100 = *p*<0.001). (B) The same results were seen when measuring nerve conduction velocity. Statistical significance was analyzed using an unpaired t-test. Nine P35 and five P>100 animals of each condition were tested (n=9 at P35 and n=5 P>100).

### Correlation of non-myelinating SCs and macrophages

Next, we investigated macrophage abundance in the sciatic nerves of WT and loSMA mice by immunostaining for F4/80, a typical macrophage surface marker. Macrophages can be recruited by SCs upon nerve damage and correlation could link SC dysfunction to macrophage recruitment in loSMA ^20^. An increase in F4/80^+^ macrophages was detected at P35 in loSMA mice (p<0.05) but not at any other timepoint. To further investigate the possible interaction between macrophages and non-myelinating SC, numbers of F4/80^+^ cells and Sox2^+^ cells were correlated. Interestingly, at P20 only WT mice showed a significant correlation (p=0.02). Meanwhile, at P35, a significant correlation was evident only in loSMA mice (p=0.005), which remained present at P70, though no longer statistically significant (p=0.09). At P>100, both WT (*p*=0.002) and loSMA (*p*=0.007) mice exhibited significant correlation (Fig. 6).

**Fig. 6.**
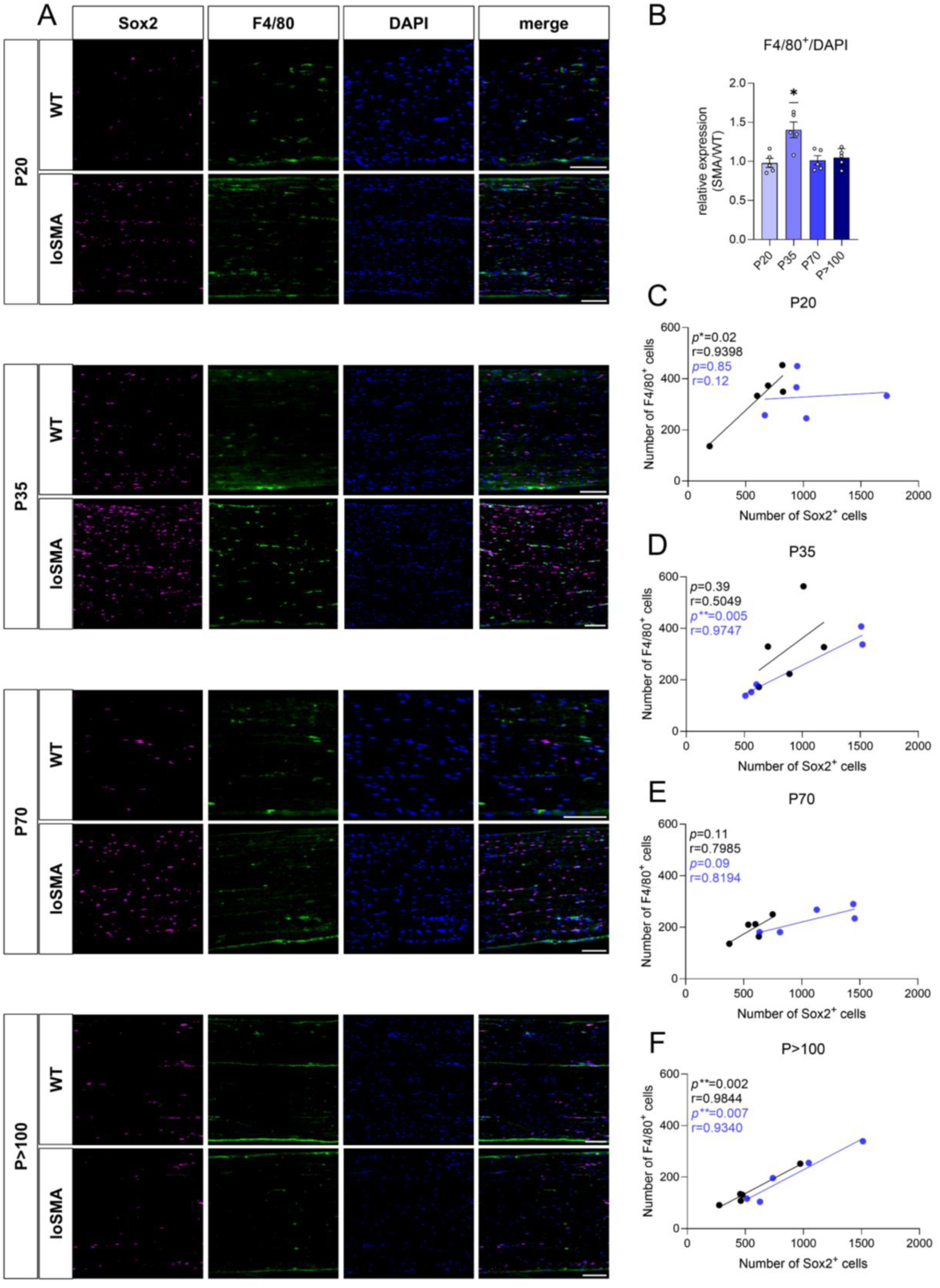
Abundance of F4/80+ cells and correlation to Sox2^+^ SC. (A) Sciatic nerves of WT and loSMA mice were cut longitudinally and immunostained for macrophage marker F4/80 (green), SC marker Sox2 (magenta) and DAPI (blue), images show representative examples of each time point. Merged images show an overlay of all channels. (B) The number of F4/80^+^ cells in relation to DAPI signals was significantly elevated at P35 (*p*<0.05), but not at the other time points. (C) The number of F4/80^+^ cells and Sox2^+^ cells showed a significant correlation at P20 in WT mice (*p*=0.02) but not in loSMA mice. (D) However, at P35, the correlation was significant for loSMA mice (*p*=0.005) but not WT mice. (E) At P70, there was no longer any statistical significance, but loSMA still showed a correlation at *p*=0.09. (F) At P>100, both WT (*p*=0.002) and loSMA (*p*=0.007) mice exhibited significant correlation. * = *p*<0.05, statistical significance for (B) was analyzed using a one-sample ratio t-test by testing the logarithmic quotient of SMA/WT against a hypothetical value of 1. For (C) to (F) simple linear regression was calculated. Five animals of each condition and age were tested (n=5). Scale bar equals 100 µm.

## Discussion

This study identifies age-dependent peripheral nerve abnormalities in a mild mouse model of loSMA. The four-copy *SMN2* model progresses through an early stage preceding overt motor neuron loss, an onset stage around P35, and later stages characterized by established motor dysfunction ^13, 15^. This temporal framework allowed SC-associated changes to be positioned relative to disease progression. The most consistent pattern emerged from P35 onward and comprised increased g-ratios, increased Sox2^+^ SC proportions, transient macrophage accumulation at P35, and later reductions in MBP immunofluorescence and electrophysiological function. These findings extend the concept of glial involvement in loSMA from central astrocytes to the peripheral nerve and suggest that peripheral remodelling accompanies the slowly progressive disease course. Studies in severe early-onset SMA models have previously demonstrated SC abnormalities, but the present data indicate that related changes also occur in a milder, later-onset context ^16, 17^.

Sox10 is a transcriptional factor found in all SCs while Sox2 inhibits myelination and is therefore present in non-myelinating SCs ^20, 21^. The increased Sox2^+^/Sox10^+^ population in loSMA nerves may therefore reflect expansion of non-myelinating SCs, acquisition of a repair-like phenotype, or incomplete SC differentiation. Further experiments will have to distinguish these possibilities. The time point P35 points towards dysfunction of SCs streamlined with astrocytes in the CNS. Dedifferentiated Repair SCs can act as pro-inflammatory regulators upon trauma ^22^. These cells can recruit macrophages as local immune effector cells into the nerve to facilitate debris clearance, stimulate an immune response, or block myelination ^20^. This provided the rationale for examining F4/80^+^ cells. Their transient increase at P35, together with the association between F4/80^+^ and Sox2^+^ cell numbers, is consistent with coordinated SC-macrophage responses at disease onset. Because the analysis is cross-sectional and based on a small number of animals, it does not demonstrate SC-mediated macrophage recruitment. Nevertheless, this indicates a dysregulation of SCs at this crucial time point before MN loss.

The correlations observed at P20 and P>100 may reflect different biological contexts. Resident macrophages contribute to peripheral nerve development, tissue surveillance, and homeostasis ^23–25^, whereas macrophage abundance can increase with peripheral nerve aging ^26^. The correlation detected in WT but not loSMA mice at P20 could therefore indicate altered developmental coordination, although this interpretation remains speculative. At P>100, correlations in both genotypes may partly reflect age-associated macrophage accumulation. More broadly, innate immune dysregulation has been reported in SMA models ^27, 28^, but cell-specific inflammatory profiling will be required to determine the phenotype and function of the macrophages observed here.

The increase in Sox2^+^ SCs may also have clinical relevance beyond motor myelination. Non-myelinating SCs support small-caliber sensory axons, and subclinical small-fiber abnormalities have been demonstrated in ambulatory adults with SMA type 3 using corneal confocal microscopy. In that study, reduced corneal nerve fiber measures correlated with motor performance ^29^. The present data do not establish small-fiber neuropathy or identify the Sox2^+^ cells as Remak SCs; nevertheless, they support further investigation of small-fiber and non-myelinating SC pathology in loSMA.

Analysis of sciatic nerve cross sections revealed deficits in myelin sheaths of loSMA mice. Axonal diameter is an important factor in myelination as only axons with a diameter of at least 1 µm are myelinated by SCs ^30^. Axonal Neuregulin-1 type III (NRG1 type III) is a key determinant of SC ensheathment and myelin thickness. NRG1 type III levels are decreased in spinal cord and ventral root axons of early-onset SMA mice and elevation leads to an improvement in axon diameter and myelination ^31^. However, whether the myelin abnormalities observed here result from altered NRG1 signalling, reduced axonal caliber, or intrinsic SC dysfunction remains to be determined. Reduction of axon diameter in loSMA mice may indicate shrinkage due to axonal atrophy, hampered axonal development or radial growth as already shown in early-onset SMA mice ^32^. This could be linked to the ubiquitous loss of SMN protein in all neurons and resulting defects in the cytoskeletal organization ^33^. Enlargement of axons, as seen in P>100 loSMA mice, could be a compensatory sprouting mechanism or the result of pathological swelling due to transport-related buildup ^34, 35^.

In accordance with abnormal axon diameter, myelin sheaths were changed in loSMA mice as well. While g-ratio measurements of the soleus nerves of early-onset SMA mice showed no difference, measurements of the intercostal nerves of early-onset SMA mice are elevated ^16, 36^. An increase in g-ratio across all time points, with simultaneous thinning of myelin sheaths in loSMA, points to defective myelination of SCs, the severity of which may depend on the mouse model used and the nerve examined. The g-ratio is a parameter for sufficient myelination of peripheral nerves, and its elevation correlates with defects in myelination capacity of SCs in various animal models with SC- and myelin-related mutations ^36, 37^. Interestingly, at P20, loSMA mice exhibited a lower g-ratio than WT mice, which may reflect a compensatory mechanism by SCs to compensate for restricted functionality. The impairment of glial cells at P35 could be too extensive to be compensated by hypermyelination, resulting in an elevated g-ratio and decreased myelin thickness from this time point onwards. Furthermore, axonal distribution is altered at P35, although not significantly so, similar to that in early-onset SMA mice ^38^. The tendency toward an increased number of axons per area suggests dysregulated axonal distribution and size.

SCs possess a high plasticity, making them responsive to changes in their surroundings and enabling them to exert compensatory mechanism to maintain nerve function ^39^. Even though myelination appears defective from a young age in loSMA mice, electrophysiological properties are affected only in older individuals (P>100). Both the slower nerve conduction velocity and the reduced amplitude suggest that compensatory mechanisms may be failing as the disease progresses. In accordance with that, MBP is also only lowered in old age loSMA mice while MBP levels were found to be increased in sciatic nerves of early-onset SMA mice ^16^. Still, CMAP reduction could also reflect MN loss, neuromuscular junction dysfunction, or muscle atrophy.

Some limitations in interpreting the results remain. The present study cannot distinguish primary SC dysfunction from secondary responses to chronic motor axon pathology. SC-specific genetic manipulation, isolated-cell experiments or controlled axon–SC co-cultures will be required to resolve this question.

Nevertheless, the findings of this study suggest that SC pathology is present in loSMA mice, making it a promising target for future therapeutic strategies. Even though restoration of SMN in SCs alone does not rescue SMA phenotypes in mouse models it improves the outcome of the mice and highlights the importance of SCs in SMA pathogenesis ^17^. SMN-directed therapies can stabilize or improve motor function in adolescents and adults with loSMA, responses are heterogeneous, and established neuromuscular deficits may not be fully reversible. Adjunctive strategies targeting downstream or non-neuronal disease mechanisms may therefore complement, rather than replace, SMN restoration ^4, 40, 41^. Treating other glial cell types, such as astrocytes, without altering SMN levels has yielded promising outcomes in mouse models^13, 14, 42^. A combination of SMN-enhancing drugs and targeting different glial cells to preserve neurons and glial homeostasis, therefore, may be the future of SMA treatment.

## Conclusion

This study identifies age-dependent abnormalities in sciatic nerve myelin morphology, an increased proportion of Sox2^+^ SCs and transient macrophage accumulation in a mouse model of loSMA. These changes precede or accompany late-stage electrophysiological impairment, but their causal relationships remain unresolved. In particular, the present experiments cannot determine whether the SC-associated phenotype is cell autonomous or secondary to chronic axonal dysfunction. Future studies will be required to establish whether peripheral glial mechanisms represent tractable adjunctive therapeutic targets.

## Author contributions

ML conceptualized and designed the study. KCL, NB, and SH performed experiments. KCL, NB, and LIS performed data analysis. KCL, LIS, and ML designed figures and the graphical abstract. LIS, NB, SH, CK, ML, and TH helped with data interpretation. KCL wrote the manuscript. LIS, NB, SH, CK, ML and TH reviewed the manuscript. ML and TH supervised the study. All authors read and approved the manuscript.

## Acknowledgements

We would like to thank former lab member Kristina Wagner for her support in sample preparation. We also thank the Imaging Center Essen (IMCES) at the Faculty of the University of Duisburg-Essen, Germany for providing access to the Leica EM UC7 ultramicrotome which is funded by the Deutsche Forschungsgemeinschaft (DFG, German Research Foundation) - 274299030, INST 58219/40-1 FUGG.

